# Multimodal electrocorticogram active electrode array based on zinc oxide-thin film transistors

**DOI:** 10.1101/2022.07.22.500376

**Authors:** Fan Zhang, Luxi Zhang, Jie Xia, Wanpeng Zhao, Shurong Dong, Zhi Ye, Gang Pan, Jikui Luo, Shaomin Zhang

## Abstract

Active electrocorticogram (ECoG) electrodes can amplify the weak electrophysiological signals and improve the anti-interference ability, but the traditional active electrodes are so opaque that cannot realize photoelectric collaborative observation. Here an active and fully-transparent ECoG array based on zinc oxide-thin film transistors (ZnO-TFTs) was developed as the local neural signal amplifier for electrophysiological monitoring. The transparency of the proposed ECoG array was up to 85% which is superior to previous reported active electrode array. Various electrical characterizations demonstrated its ability of electrophysiological signal recording, with a higher signal-to-noise ratio of 19.9 dB compared to the Au grid one (13.2 dB). The high transparency of ZnO-TFT electrode array allowed the collecting electrophysiological signals under direct light stimulation on optogenetic mice brain concurrently. The ECoG array could also work under 7-Tesla magnetic resonance imaging to record local brain signal without affecting brain tissue imaging. As the most transparent active ECoG array to date, it provides a powerful multimodal tool for brain observation, including recording brain activity under synchronized optical modulation and 7-Tesla magnetic resonance imaging.

## Introduction

Electrocorticography (ECoG) is a technique that collects electrophysiological signals from cerebral cortex through a planar electrode array above or below the dura mater.^1^ ECoG signals have been used to monitor and locate the neurological dysfunctions such as intractable epilepsy.^2^ In the past few years, as a new emerging technology, brain-computer interface (BCI) combining with ECoG allows the development of closed-loop systems that achieve neuroprosthetic applications for motor functional reconstruction,^3^ and decoding of vision, ^4^ movements^5^ and speech.^6^ Various micro electrode arrays have been developed to enable highly reliable neural mapping in the cerebral cortex with high spatiotemporal resolution with no damage to brain tissues.^7,8^

For wired ECoG electrodes, neural signals are transmitted through wires before being amplified and processed by external circuits, which makes them extremely sensitive to environmental noise.^9^ Thus, active electrode arrays^10^ have been studied and developed to improve signal-to-noise ratio (SNR) through on-chip pre-amplifiers that mainly consist of transistors. For active devices, the neuronal transmembrane current conducted through the electrolyte will polarize the gate electrodes and change the conductivity of the active channels, which forms local amplifiers.^11^ Several types of transistors, such as the solution-gated field effect transistor (SGFET),^12^ organic electrochemical transistor (OECT)^11,13^ and ion-gated organic electrochemical transistor (e-IGT),^14^ have been adopted to monitor neural activity, indicating the advantages of the transistor-based active electrode arrays in terms of low noise level.

With the development of optical neuromodulation and optical imaging techniques in neuroscience, the demand for optically transparent electrode arrays is rising rapidly. Transparent electrode array is able to provide an interface which is feasible for simultaneous and in-situ optical modulation and electrical monitoring. Therefore, active electrode arrays with high transparency, including transparent electrodes and transparent active circuits consisted of amplifiers and interconnects, are highly required. Although some conductive transparent materials in passive electrode arrays, such as the ultra-thin metal films with nanometer thickness,^15,16^ indium-tin-oxide (ITO),^17,18^ graphene, ^19,20^ PEDOT:PSS,^21^ etc. offer possible solutions to the transparent electrode arrays, until now, there is no highly transparent active electrode array because of some difficulties discussed below.

The first difficulty is that the small spontaneous electrophysiological potential of neurons (tens to hundreds of micro-volts) and photoexcited potentials (tens to thousands of microvolts) are easily buried in the light-induced artifacts caused by photosensitive transparent materials.^22,23^ Therefore, electrical and optical properties of the devices need to be stable under light stimulation of different wavelengths and powers, so that it does not affect the neuron signal recoding. Secondly, since transparency and conductivity of materials often require a compromise,^24^ complicated structural design and extensive engineering must be taken to select appropriate transparent conductive materials and combine it with transparent semiconductor materials for the device fabrication. Furthermore, the structure and processing techniques should be compatible with that for ECoG array, and the transistors produced by the same process are supposed to possess stable electrical and optical performance. Additionally, for in-vivo applications, it is necessary for the transistors to meet strict requirements, such as biocompatibility,^25^ long-term stability,^26^ etc., making it much more difficult to come up with an appropriate solution for transparent transistors. In 2017, Lee et al. implemented a transparent active electrode array via OECTs, achieved an transparency of 60% through gold wires.^13^ This is the only study to date on the transparent active ECoG grid, but the transparency of the electrode array is limited by the usage of ultrathin metal grids.

The high transparency of thin film transistor (TFT) makes the TFTs-based transparent active ECoG electrode arrays very feasible and attractive. TFT is a kind of FETs composed of thin-film elements on a substrate,^27^ which is usually used in large-area electronic products, such as liquid crystal displays,^28^ light-emitting displays.^29^ Some transparent conducting oxides (TCOs) materials, such as ITO,^30^ ZnO,^31^ and cadmium-oxide,^32^ etc., have been incorporated to enable fully transparent TFTs. ZnO is considered the most suitable material for active channel layer material in TFT, as it has excellent semiconductor characteristics (high field effect mobility) and high optical transparency (insensitivity to visible light owing to its wide bandgap).^33^ This inspires us a promising solution for highly transparent active electrode array for neural electrophysiology.

In this work, a transparent active electrode array for neural signal recording on brain surface with ZnO-TFTs as the pre-amplifiers (we name it ZnO-TFTs electrode array) is firstly developed and investigated systematically. The ZnO-TFTs are composed of ITO conductor, aluminum oxide (Al_2_O_3_) insulator and ZnO semiconductor, and finally passivated by Al_2_O_3_ with tens of nanometers thickness. The ZnO-TFTs exhibited stable electrical properties and high optical transparency up to 85% in the visible light range. The ZnO-TFTs electrode array was then employed to record spontaneous sleep waveforms, demonstrated its feasibility for neural signal acquisition. Owing to the adoption of pre-amplifiers, the SNR of our ZnO-TFTs electrode array reached 19.9 dB, superior to that of the passive Au electrodes (13.2 dB). Owing to its high transparency, it allowed in-situ optical stimulation and simultaneous neural electrophysiological recording, to map out the distribution of light-evoked potential from an optogenetic mice brain. Furthermore, the ECoG array was applied in 7-Tesla (7T) ultra-high magnetic field, and obtained clear magnetic resonance imaging (MRI) with no blurring, demonstrated its feasibility in MRI-ECoG synchronous recording.

## Materials and methods

### ZnO-TFTs circuits design

The amplifier and address gate circuits were designed based on n-type LTPS-TFT SPICE model (level 62) and simulated using the HPICE software (Synopsys). The threshold voltage was set to be 0 V according to previous study. ^31^

### ECoG array fabrication

The processing steps for the ZnO-TFTs electrode array is shown in Figure S1. The layout of the ZnO-TFTs electrode array was drawn using Tanner EDA software, as shown in Figure S2. The ZnO-TFTs circuits and electrode array were integrated on the same substrate by the multi-layer process discussed in Supporting Information. Thin film ITO was employed in our ZnO-TFTs, since it has been commonly used in transparent electrode arrays because of its high transmittance in the visible spectrum, high electrical conductivity,^34^ and biocompatibility.^35^ And an ALD Al_2_O_3_ film was used for encapsulation layer. It is because Al_2_O_3_ is considered as one of the most biocompatible materials,^36^ while ALD Al_2_O_3_ film was pinhole-free films, which is an excellent electrical insulator and moisture barrier.^37–39^ Finally, the ZnO-TFTs array was bonded with flexible printed circuit as described in Supporting Information.

### Characterization

Electrical properties of the devices were characterized by a semiconductor characterization system (Keysight B1500A) on a probe station (Figure S3). Transfer characteristics of the ZnO-TFTs were assessed by applying a drain voltage (*V*_*D*_) of 5 V and a gate voltage (*V*_*G*_) between -10 V and 15 V, and the transconductance was calculated by differential method. The output characteristic curve was obtained for *V*_*D*_ between 0 V and 10 V and *V*_*G*_ between -1 V and 5 V with a step of 1 V. Mechanical flexibility was assessed under a curvature of 15 cm in radius, and the transfer characteristics were measured after 0 cycles and 10 cycles.The transparency of the ZnO-TFTs array was evaluated by an UV-Vis spectrophotometer (Cary 60 Series, Agilent Technologies, Inc.) in the wavelength range from 200 nm to 800 nm.

### In vitro experiments

For all subsequent measurements, the bias voltages of all the ZnO-TFTs electrode array were set to *V*_*D*_ = 2.5 V and *V*_*S*_ = -2.5 V respectively, unless specified. Characterization for frequency response and response time were also conducted on a probe station. For frequency response tests, a probe was placed on a gate electrode pad to deliver sinusoid waves with a peak-to-peak value (*V*_*p−p*_) of 1 mV at a frequency in the range from 0.1 to 300 Hz generated by a weak electrophysiological signal simulator (SKX-8000 Series, Ming Sheng Electronic Technology Co., Ltd, Xuzhou, China). To characterize the response time, a square wave of 40-ms duration, 0.5-mV amplitude and 50% duty cycle was generated by the simulator and applied to the gate electrode, and the corresponding output signals were recorded with a sampling rate of 16 kHz. For electrical characterization in saline, a dupont line transferred the sinusoid wave (25 Hz, 2-mV *V*_*p−p*_) generated by the signal simulator to saline solution (0.9% wt) in a beaker. Meanwhile, a ZnO-TFTs electrode array was submerged in the saline, so that the gate electrode could collect signal transmitted through the saline solution. The output signal from the ZnO-TFTs was transfered by an I-V converting circuit and then recorded by a customized electrophysiological signal recording system.

### Animals

SpragueDawley rats (250-300 g) and C57BL mice (25 g) were used. All surgical and experimental procedures conformed to the Guide for The Care and Use of Laboratory Animals (China Ministry of Health) and were approved by the Animal Care Committee of Zhejiang University, China. Animals were fully accustomed to environment for 7 days before any experimental procedure and were given standard rat chow and water.

### Electrophysiological recording on rat brain surface

A rat was first treated with atropine by intra-abdominal anesthesia with a dosage of 0.12 mg/100 g to relieve pain. The rat was then anesthetized with propofol (10 mg/ml solution, 1.2 ml/100 g) via intraperitoneal injection (IP) according to its weight and set in stereotaxic head frame. An incision was made on the brain skin, and a dry and clean skull was exposed after H_2_O_2_ treatment. A cranial nail was screwed into posterior fontanelle skull as ground. A cranial window was drilled to expose a 7 *×* 5 *mm*^2^ window on right hemisphere. Mannitol was injected by IP to prevent edema. The ZnO-TFTs electrode array was fixed on the arm of dorso-ventral axis, and implanted position was carefully adjusted by micro-drive screw of three-dimensional axis. Neural signal was recorded by customized recording system mentioned above. After data acquisition, the ZnO-TFTs electrode array was removed, and an Au electrode array was implanted into the same position for neural signal recording. The rat was then euthanized. After rat died, the ZnO-TFTs electrode array and Au electrodes were implanted to the same region, and the noise was recorded by two arrays separately.

### Optogenetics

AVV-hSyn-hChR2(H134R)-mCherry virus was injected to primary somatosensory cortex (S1, AP: -1.2 mm, ML: +2.2 mm) of a male mouse. Two weeks were needed for the mice to recover from the surgery and for viral expression. For optogenetics, the mouse was anesthetized, and set in the stereotaxic head frame. A cranial window was conducted around the virus injected site using an electrical cranial drill, and the dura was removed. ZnO-TFTs electrode array was implanted to the cranial window, with the gate electrode positioned precisely in the virus injected region. And the ground wire was connected to a cranial nail fixed on the posterior fontanelle. A blue laser of 473 nm wave length (Shanghai Laser & Optics Century Co., Ltd., China) was used to illuminate the gate electrode region. Each light pulse train was triggered by an electrical stimulator (Master-8, A.M.P.I Instruments, Ltd., Israel), it had 30 trials at 4 Hz, with a pulse width varied from 3 ms to 20 ms and an excitation power from 1mW, 5 mW, 13 mW and 40 mW.

### Light-induced artifacts

A ZnO-TFT electrode array was placed on agarose and the reference/ground wire was inserted into the agarose. Laser stimulation with different pulse widths and powers shone on the gate electrode region, and the evoked potential was measured by recording system.

### Data acquisition and processing

All recordings were conducted in the unipolar mode at a sampling frequency of 1 kHz unless specified, and filtered with a high-pass filter of 0.2 Hz (4 orders, Butterworth) and a notch filter of 50 Hz in MATLAB (MathWorks). For optogenetics and light-induced artifacts, the evoked signals of 30 trials were averaged for the assessment.

### MRI imaging

A piece of ZnO-TFTs array was put on the surface of the rat cadaver brain, across the left and right hemispheres. The dura of the rat brain was removed and saline was applied to keep the brain hydrated. MRI scanning was acquired using a 7T research scanner (Magnetom, Siemens Healthcare, Erlangen, Germany). The rat brain was placed in the head coil of the 7T MRI scanner. The sequence had TR of 410 ms, TE of 7.56 ms and slice thickness of 1 mm.

## Results and discussion

### Structure of ZnO-TFTs electrode array

To enable the neural interface with high-transparency and high signal-to-noise ratio, we developed an active, non-penetrating electrodes based on the transparent ZnO-TFTs. The structure of ZnO-TFT is shown in Figure 1A. The ZnO-TFTs electrode array was fabricated using a multi-layer process as discussed in Materials and Methods.

**Figure 1.**
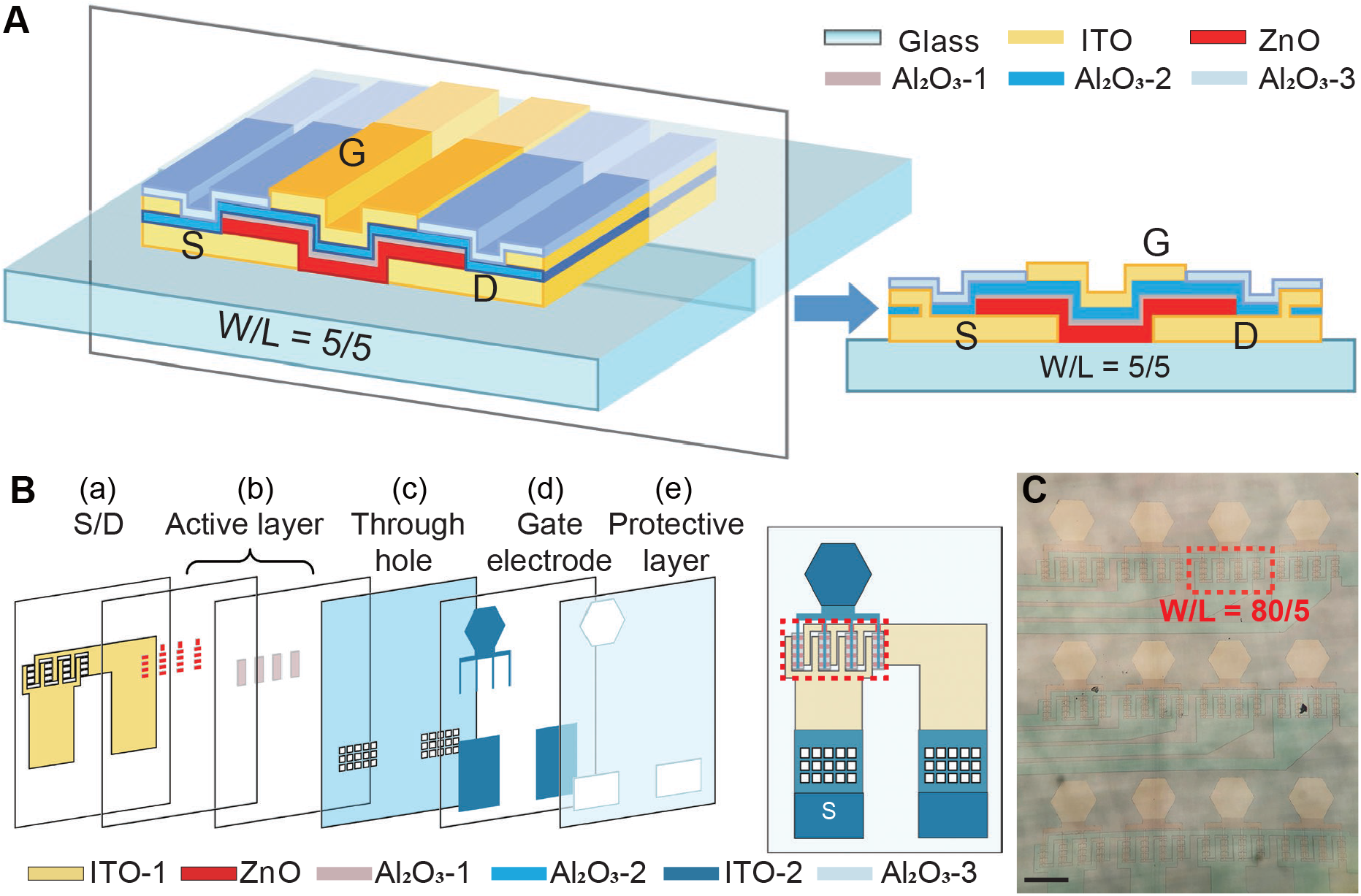
Schematic illustration and layout of the transparent ZnO-TFTs array. (A) Three-dimensional structure diagram (left) and the cross-section (right) of the transparent ZnO TFT. (B) The exploded view (left) and structure (right) of a ZnO-TFTs electrode. (C) Microscope image of a 3 *×* 4 ZnO-TFTs array. The red frame labels an active region consisting of 16 paralleled ZnO-TFTs, same as that in the red frame of Figure 1B (right). Scale bar: 200 µm.

The structure diagram of a ZnO-TFTs electrode is shown in Figure 1B. The size of ZnO-TFTs array is 10.8 *×* 7.0 *mm*^2^, and each array consists of 3 *×* 4 electrodes with 350-µm-spacing horizontally and 620-µm-spacing vertically, as shown in Figure S2. A photograph of an ZnO-TFTs electrode array is shown in Figure 1C.

### Design and characterization of ZnO-TFTs array

Simulations were conducted to optimize performance of the ZnO-TFTs as the pre-amplifer. We set the width to length ratio (W/L) to 5/5, 10/5, 20/5, 40/5 and 80/5 respectively for the TFTs in the simulation. The calculated transfer characteristics and transconductance *g*_*m*_ are shown in Figure S4A and Figure S4B respectively. It shows that once the quiescent operating point is determined, the transconductance has a linear relationship with W/L in the linear region and then saturates with the increase of W/L ratios. We fabricated ZnO-TFTs with different W/L, and characterized their electrical properties. As expected, the transfer characteristics of the fabricated ZnO-TFTs with larger W/L ratios has larger linear slopes (Figure 2A), which give larger *g*_*m*_. By deuterium plasma treatment, the threshold voltage of the transistors reduced than the simulation, causing the transistors to exhibit typical depletion mode, as shown in Figure 2A. The disagreement between the experimental results with the simulation ones is mainly attributed to the deuterium plasma treatment. Finally, to obtain a better amplifying capability, ZnO-TFTs with a W/L ratio of 80 µm/5 µm were selected to be the ECoG array for the following experiments. It is worth noting that each transistor array is consisted of 16 paralleled transistors through symmetrical layout technique (red frame labeled as shown in Figure 1C and the top-right inset of Figure S2), to avert the systematic mismatch caused by process.^40^

**Figure 2.**
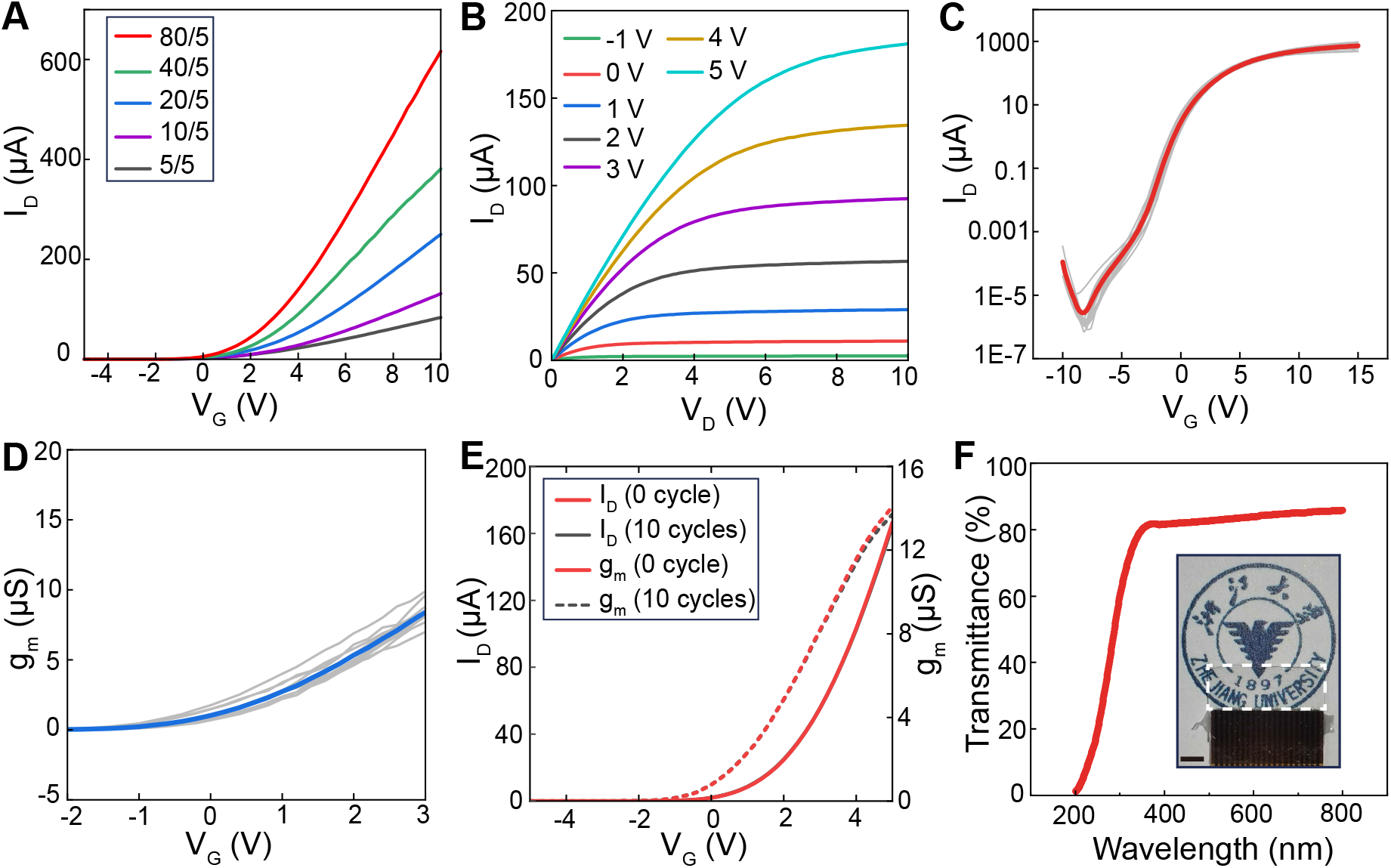
Electrical, mechanical and optical properties of the ZnO-TFTs array. (A) Transfer curves of the TFTs with different W/L ratios at *V*_*D*_ = 5V. (B) Output characteristics of TFT with W/L = 80/5, showing the drain current (*I*_*D*_) as a function of *V*_*D*_ with *V*_*G*_ varying from -1 V to 5 V (step = 1 V). (C) Transfer characteristics of twelve ZnO-TFTs electrodes of an array. Red line is the average value. (D) The resulting transconductance of twelve ZnO-TFTs electrodes from the same array. Blue line is the average value. (E) ZnO-TFTs electrode maintained stable electrical properties after 10 bending cycles at a bending radius of 15 cm. (F) Transmittance spectrum of the ZnO-TFTs array. Inset is an optical image of 3 *×* 4 ZnO-TFTs array, showing its high transparency. The white frame labels the electrode array. Scale bar: 2 mm.

The output characteristics of the fabricated ZnO-TFTs with W/L=80/5 showed good performance in both the linear and saturation regions (Figure 2B), which is consistent with the simulation results (Figure S4C). The transfer characteristic curves and corresponding transconductance curves of twelve electrodes in one ZnO-TFTs array are illustrated in Figure 2C and D respectively, demonstrated good consistency among transistors on the same piece. According to the simulation, the transconductance reaches the maximum value when *V*_*G*_ is about 4.5 V (Figure S4B). However, from the measurements, we found that when *V*_*G*_ was increased above 3 V, the consistency of *g*_*m*_ among different electrodes was relatively poor. Therefore, as a compromise, in the subsequent experiments we choose the operating point of *V*_*G*_ = 2.5 V and drain-source voltage (*V*_*D*_) = 5 V, respectively, where *g*_*m*_ is 6.8 µS. Notably, the steady state gate current is always less than 0.35 nA, for *V*_*G*_ between -10 and 15V, and *V*_*D*_ of 5V (Figure S5).

We bended the ZnO-TFTs array to a 15 cm radius of curvature for 10 times, and measured the transfer characteristics of active electrodes before and after bending (Figure 2E). The transfer characteristic curve is not affected visibly by bending, demonstrated the flexibility of our ZnO-TFTs array. The average optical transmittance of the device (including the substrate of glass) in the visible spectrum (380 to 780 nm) is large than 80% (Figure 2F), enabling the device to combine optical stimulation, optical imaging and electrophysiological recording together for the experiments. In the inset of Figure 2F, an optical image of a ZnO-TFTs electrode array distinctly demonstrates its high transparency.

### In vitro evaluation

As shown in Figure 3A, neural signals could be collected by the ZnO-TFTs electrodes, whereas the alternating voltage difference (*V*_*e*_) between the gate and source could control the alternating current of the ZnO-TFTs (*i*_*ds*_). The output signal is collected by a customized recording system.

**Figure 3.**
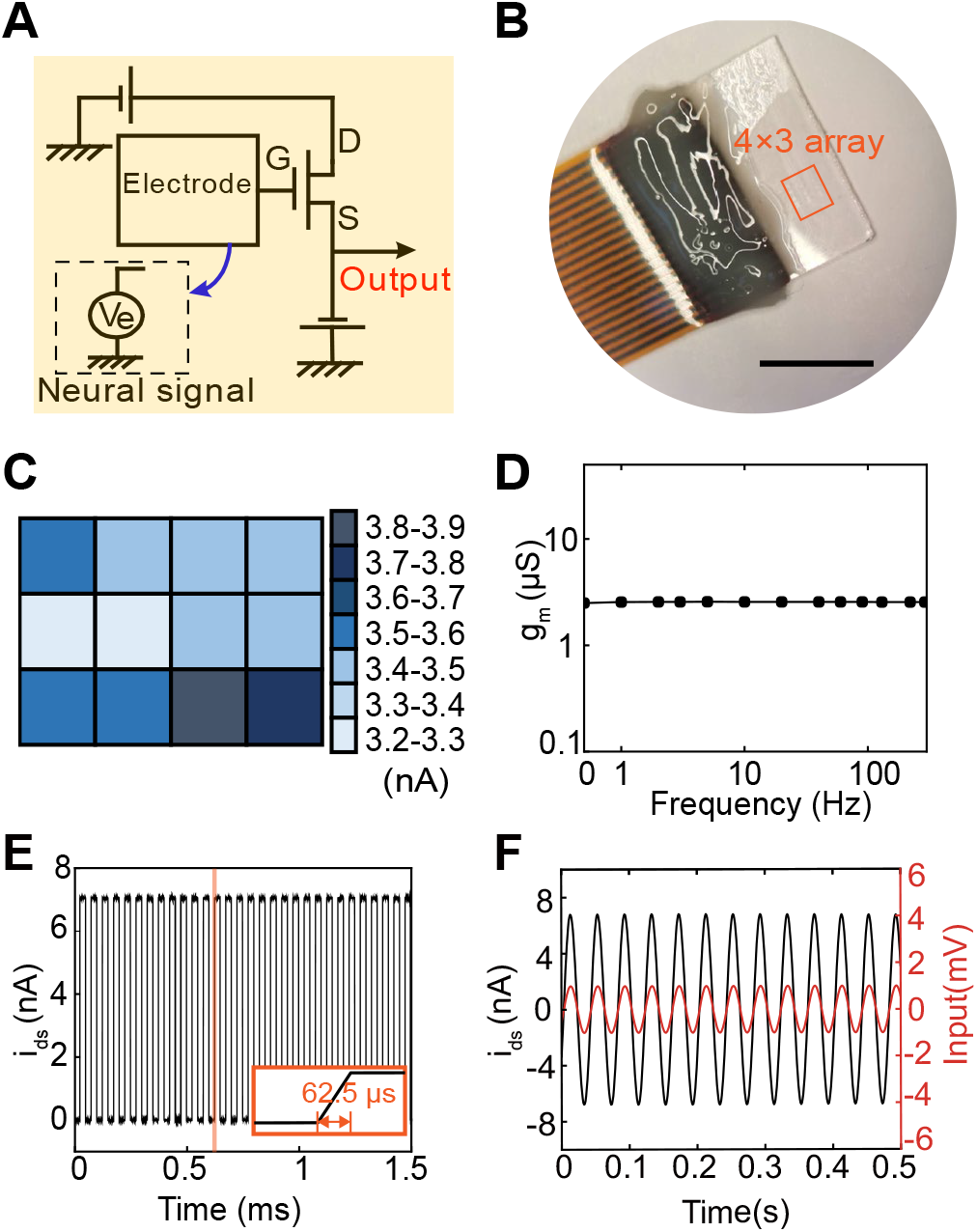
Feasibility of neural signal acquisition from the ZnO-TFTs array. (A) Wiring layout of the ZnO-TFTs electrode array. Ve represents the electrophysiological signals. (B) An image of a 3 × 4 transparent ZnO-TFTs array. Scale bar: 0.5 cm. (C) Distribution of *i*_*gs*_ from the 12 ZnO-TFTs with a sine wave input at a frequency of 60 Hz, and 1-mV *V*_*p−p*_. (D) Frequency response characteristics of the ZnO-TFTs electrode array. (E) Response characteristics of a ZnO-TFTs electrode. The response time is less than 62.5 µs. (F) Sine wave recorded by the transparent ZnO-TFTs electrode in saline solution with an input of a 25-Hz frequency, 2-mV *V*_*p−p*_ sine wave.

When a 60-Hz, 1-mV *V*_*p−p*_ sinusoid wave is transmitted to the 12 electrodes of a 3 *×* 4 ZnO-TFTs array (Figure 3B), the amplitude of each electrode can be obtained as displayed in Figure 3C. The amplitudes of the ZnO-TFTs are in the range of 3.4 to 3.6 nA, indicating that the ZnO-TFTs electrodes have good consistency. The ZnO-TFTs electrodes has a flat frequency response characteristic curve (Figure 3D), which means that the ZnO-TFTs electrodes are suitable for mapping neural signal with complex frequency components. Figure S6 exhibits a typical recording for an input of a 60-Hz, 1-mV *V*_*p−p*_ sine wave. Its SNR was calculated to be 16.8 dB (2.50 nA / 0.36 nA, RMS). In addition, the latency time was measured by sending a rectangular wave to the gate electrodes and recording the output. Figure 3E shows that the delay time of the ZnO-TFTs electrodes is less than 62.5 µs (temporal resolution at 16 kHz ampling rate), indicating that the ZnO-TFTs electrodes could record neural signals at 16 kHz, meeting the requirements of neural electrophysiological recording.

The stability of electrical performance of the ZnO-TFTs in liquid or ionic solutions is important for the proposed application. Based on the previous study that summarized the failure time of ALD deposited ultra-thin Al_2_O_3_ films with different thickness in water, ^41^ we used an ALD deposited ultra-thin Al_2_O_3_ film with a thickness of 20 nm as the encapsulation layer to protect the ZnO transistors. The input sine wave and recorded signal are shown in Figure 3F, when the electrodes were completely soaked in saline. The signal recorded by ZnO-TFTs electrodes is in phase with that of the input signal, and the SNR is 19.5 dB (RMS noise level of 0.51 nA and RMS amplitude of signal of 4.81 nA). The same electrode was tested again one week later, and signal with similar SNR was obtained as shown in Figure S7. It demonstrates that the ZnO-TFT electrodes immersed into ionic liquid work well with stable electrical properties.

Table 1 summarizes the types of transistors used in varied neural electrophysiological interfaces and their optical and electrical characteristics. Different transistors such as silicon nanomembrane transistors,^42^ OECTs^11^ and Solution-Gated FETs,^12^ have been used in active ECoG arrays. However, due to the opacity of conductive materials and substrate, these active ECoG arrays was non-transparent. The transparent OECTs implements the only active transparent ECoG array, but the transparency is limited to less than 60% due to the use of metal grids, whereas the adoption of ITO in our work ensures the high optical transmittance of over 80% in the visible range. The higher transparency of ZnO-TFTs brings more advantages in optical imaging of the brain, such as single photon imaging or two photon imaging, because the higher transparency makes it possible for fluorescence to pass through the array and be observed. And ZnO-TFTs electrode array has a higher cutoff frequency than that of transparent OECTs array (50 Hz). Unlike the OECTs array, which can measure only a very limited range of low-frequency signals, ZnO-TFTs electrode array can measure electrophysiological signals with complex frequency components. Furthermore, the low latency time of our ZnO-TFTs (*<* 62.5 µs) is less than that of the transparent OECTs array (97 µs). For transparent active ECoG array, although the transconductance of our ZnO-TFTs is about three orders of magnitude smaller than that of the transparent OECTs based on PEDOT:PSS (2.2 mS maximum),^13^ the SNR of ZnO-TFTs electrodes is comparable to that of the OECTs, which is attributed to the relatively low gate-current of the ZnO-TFTs. Besides, unlike the OECTs for bioelectronics, which faces the challenge of electrical performance instability during long-term storage,^43^ the long-term electrical stability of the ZnO-TFTs in storage has been confirmed.^33^ The electrical properties of our ZnO-TFTs electrode array stored at 15 °C and 6% relative humidity for 6 months were tested (Figure S8). The results showed that the ZnO-TFTs electrode array maintained stable electrical performance for long-time storage with no visible deterioration, demonstrated its superiority, particularly useful for the proposed application.

**Table 1.**
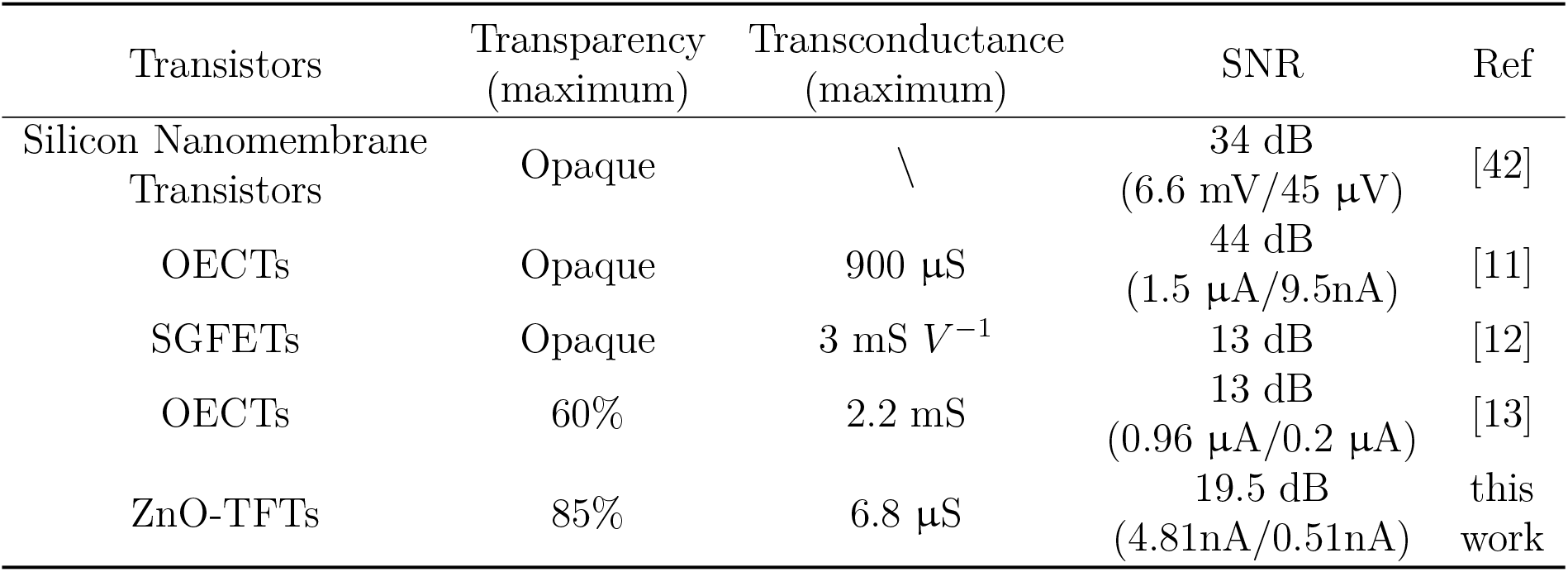
The characteristics of various active ECoG electrodes.

### In vivo electrophysiological measurements and optogenetics

The ZnO-TFTs electrode array was applied on an anesthetized rat to record brain activity. As shown in Figure 4A-B, ZnO-TFTs array and Au array were placed on the same cortical region of the open skull window for comparative experiments. Cortical electrical potentials from the two electrode arrays (Figure 4C-D) show typical episode of non-rapid eye movement sleep characterized by the high-amplitude local field potential slow (0.5-4 Hz) waves (i.e., *δ* waves).^44^ The power spectral density analysis (Figure 4E) of both the ZnO-TFTs and Au array electrodes have high power in low-frequency oscillations of the *δ* band that is associated with the sleep-like rhythms. The signals recorded after the rat died were regarded as noise. Figure S9 showed the electrophysiological recording and noise level of the two arrays. The SNR of the ZnO-TFT array electrode is 19.9 dB with a RMS amplitude of 1.69 nA for the ECoG signal and 0.17 nA for the noise, whereas the SNR of the Au electrode array is dB (70.7 µV RMS for ECoG signal, 15.5 µV RMS for noise). The ZnO-TFTs array exhibits superior SNR. These suggest that our ZnO-TFT electrode array is able to record electrophysiological signal from the cortex of anesthetized rats with high quality.

**Figure 4.**
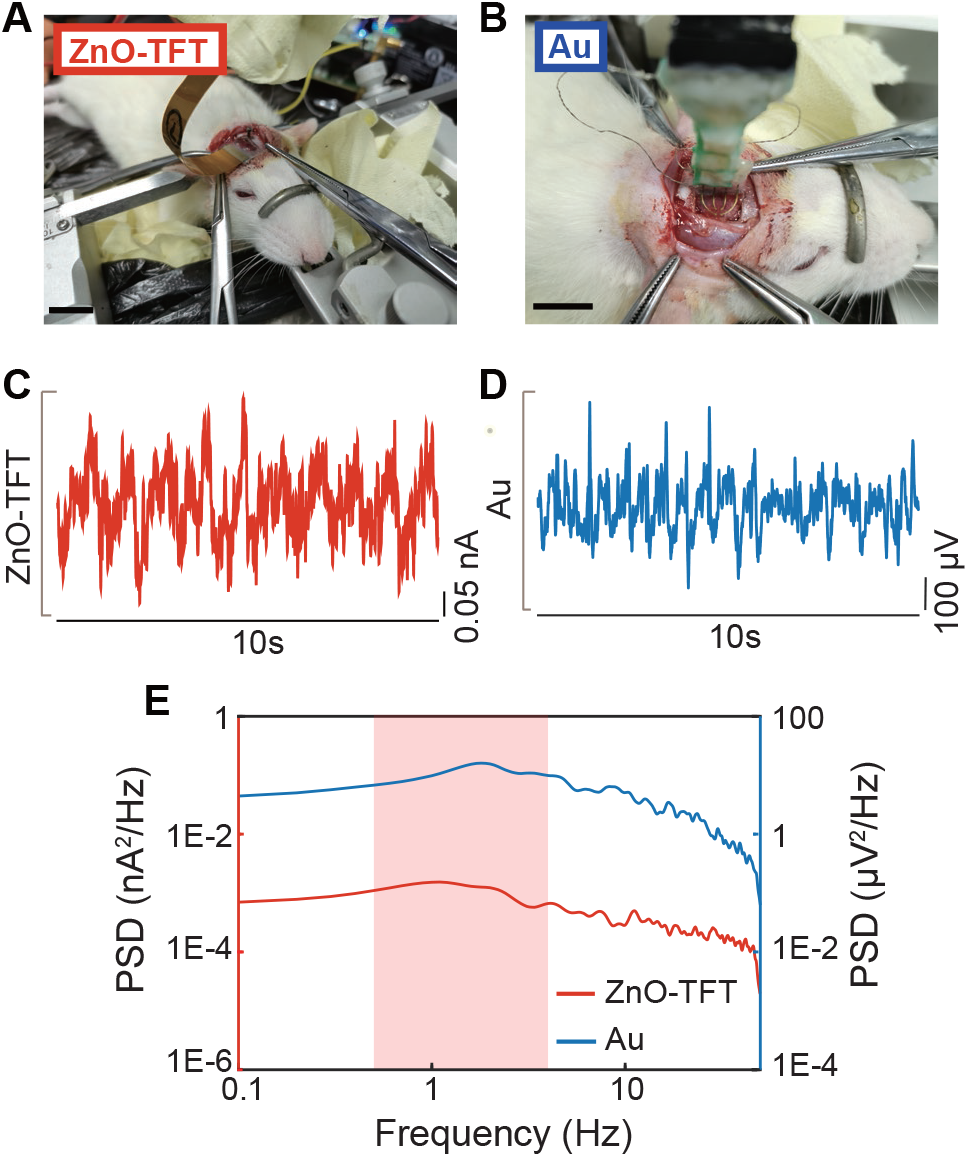
Electrical performance characterization of ZnO-TFTs array on rat brain. (A) A photo of the setup for recording sleep waveform from a ZnO-TFTs array in an anesthetized rat brain where the transparent ZnO-TFTs array is implanted. Scale bar: 1 cm. (B) A photo of the setup for recording sleep waveform from an Au array implanted to the same sites. Scale bar: 1 cm. (C-D) 10 s long trace of electrophysiological signal recorded by the ZnO-TFTs electrode (C) and by the Au electrode (D) from the anesthetized rat brain. (E) Signal power spectral density of neuronal electrical activity in (C-D).

The transparent neural electrode arrays enable light to directly modulate cortex neurons through the interface, with negligible optical power loss, making it possible to combine optogenetics with neural electrophysiology in situ and simultaneously. Therefore, for simultaneous optogenetic modulation and neural electrophysiological recording, it is necessary to evaluate the light-induced artifacts that could contaminate neural signals. Becquerel effect^22^ is considered to be the main cause of photoinduced artifacts. According to previous studies, there was no difference in the transfer characteristic of ZnO-TFTs under dark and bright (wavelength range from visible to near ultraviolet) conditions, except for a moderate hysteresis under illumination of 405 nm wavelength.^31^ ZnO-TFT devices are not as sensitive to visible light as other amorphous indium gallium zinc oxide (IGZO) TFT,^40^ mostly owing to the polycrystalline nature of the ZnO films deposited by ALD. ^33^

To further study the artifacts of ZnO-TFTs, we placed a ZnO-TFTs array electrode on agar to mimic the neural electrophysiological recording from the array on the brain surface (Figure S10). A 473 nm blue laser light was illuminated at the region of the ZnO-TFTs electrode array with the beam spot size of 1 mm. Detailed settings could be found in Methods. The amplitude of artifacts was found to depend on the stimulation light power and pulse width (Figure S11). The photoinduced artifacts of the ZnO-TFTs electrode increases sharply when the light irradiates on it and it persists for tens of milliseconds after the light is off, which is similar to the previous studies using metal electrodes.^45^ Further study will be necessary to fully characterize the overall optical-electrical properties of the ZnO-TFTs and source of potential artifacts.

Then, we performed optogenetic and neural recording using our ZnO-TFTs array. After successful expression of ChR2 virus in S1 region of mice brain, a cranial window around the injection area was drilled, and a 3 *×* 4 ZnO-TFTs electrode array was precisely placed through a stereotaxic instrument to ensure that the electrodes covered the injection site. The photograph of the implanted ZnO-TFTs array (Figure S12A) indicates that it could attached to the mice cortex, and the clear cortical vessels beneath the array also show the high transparency of the array electrode.

The activation of channelrhodopsin with a blue light stimulation would lead to the depolarization of neurons. ^46^ As shown in Figure 5A, a blue laser light of 473 nm shined on the electrode region of the ZnO-TFTs array through an optical fiber, same as the settings in artifacts measurement. Because of the high transparency of ZnO-TFTs array, the laser light could penetrate the array to activate the cortex neurons infected by ChR2 virus. Figure S12B is a photograph of the laser beam illuminated on the site on the ZnO-TFTs array laminated on the S1 cortex surface (virus injected spot) of the mouse brain. A representative recording from the ZnO-TFTs array in Figure 5B shows the evoked potentials with a larger negative amplitude generated in synchronization with the photostimulus pulses (13 mW, 5 ms, and 4 Hz). Different from light-induced artifacts, these potentials have the same frequency as that of the light pulses, similar amplitude with a long persisted tail, demonstrating their reliability as neural signals. The frequency of the light pulses is 4 Hz, which is below the threshold of ChR2 virus dynamic response. Figure 5C shows the recording of 30 trials (mean *±* std) from Figure 5B with small scattering, indicating the stability of electrical performance of ZnO-TFTs array under the same intensity of light stimulation.

**Figure 5.**
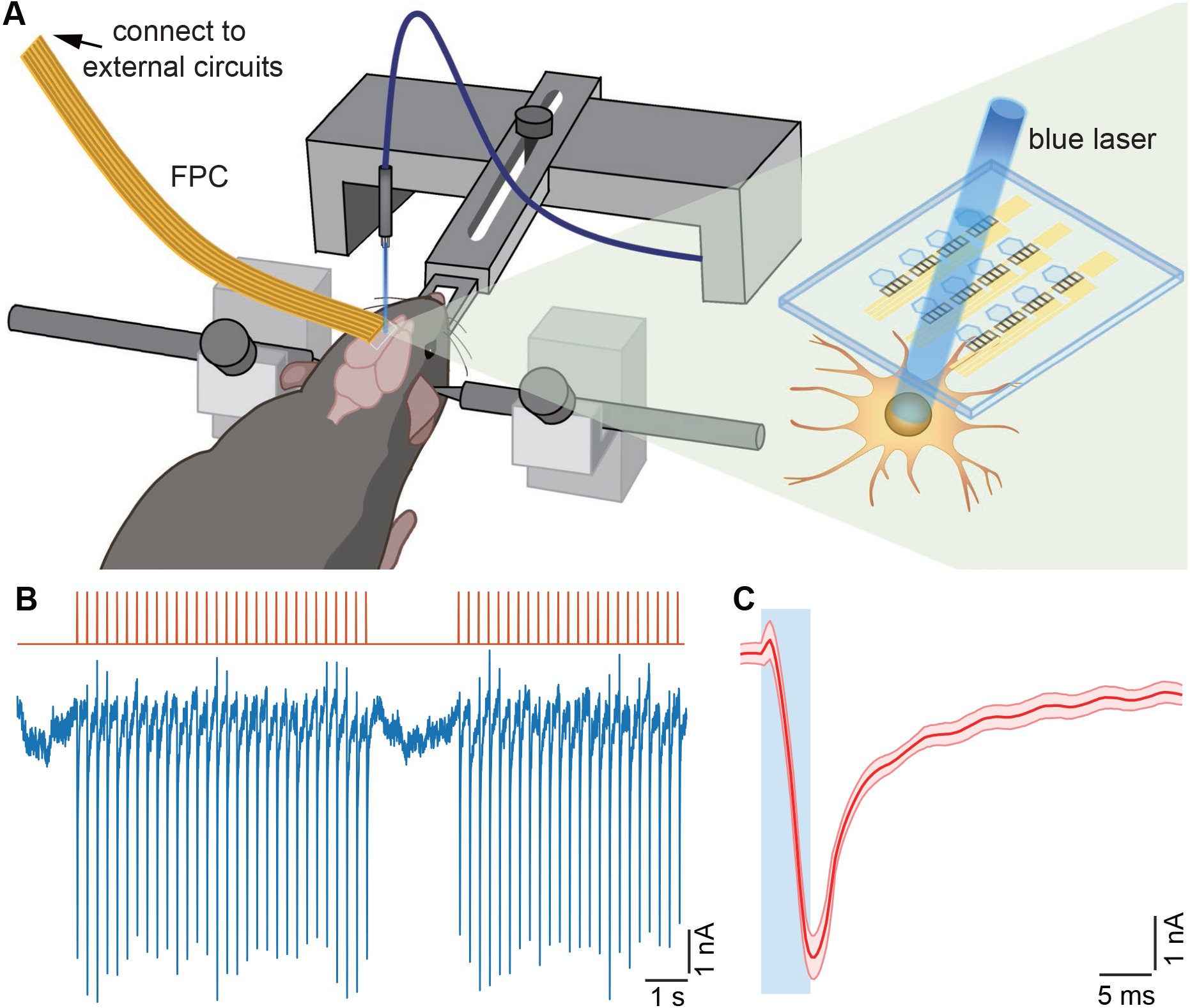
In vivo evaluation of ZnO-TFTs array by laser stimulating on ChR2-virus expressed on mouse brain. (A) Schematic of the transparent ZnO-TFTs array on S1 cortical surface (virus injection area) under 473-nm blue light pulse train stimulation transmitted by an optical fiber. (B) Top: the pulse train of the light stimulus (wavelength: 473 nm, frequency: 4 Hz, duration: 5 ms, diameter: 1 mm, and intensity: 13 mW) applied on the injection cortical surface. Bottom: neural signal potential recorded by the transparent ZnO-TFTs electrode array (row 1st, column 4th of the ZnO-TFTs array) exhibits the light-evoked negative peaks synchronized with the light stimulation train. (C) Averaged light-evoked potential (mean ± std, n = 30) of (B). The blue rectangles illustrated the start and duration of light stimulation pulses.

In addition, the effects of light pulse power and pulse width on induced neural signals were studied (Figure S13). As expected, it showed the increased neural activation in response to higher potostimulus power and wider pulse width, indicating enhanced depolarization of neurons. Specifically, higher light stimulation power would lead to faster decrease and larger negative peaks of electrical potential. And wider pulse width would result in longer depolarization periods as well as larger negative peaks of evoked neural signal.

### 7T MRI imaging on a rat brain

Many practical applications of MRI-ECoG simultaneous measurement have been introduced for the study of epilepsy, sleep, and other brain functions.^47^ Implanted ECoGs may introduce artifacts that obscure the regions of diagnostic importance, thus bring challenges to MRI imaging. The MRI imaging was conducted with the ZnO-TFTs array on a rat cadaveric model under the ultra-high magnetic field of 7T. Figure 6A exhibits the implanted ZnO-TFTs array. The array was attached tightly to surface of the rat cadaver brain, spanning the left and right hemispheres. Figure 6B, C exhibit the MRI images at two slice positions, sagittal and transversal plane respectively. The MRI images show no blur around the rat cadaver brain tissue, even in the regions close to the array, signifying that the ZnO-TFTs array has no effect on MRI imaging. Therefore, the ZnO-TFTs array could be used for MRI-ECoG simultaneous recording to study mechanisms of the brain network activity, demonstrated its great potential for the application.

**Figure 6.**
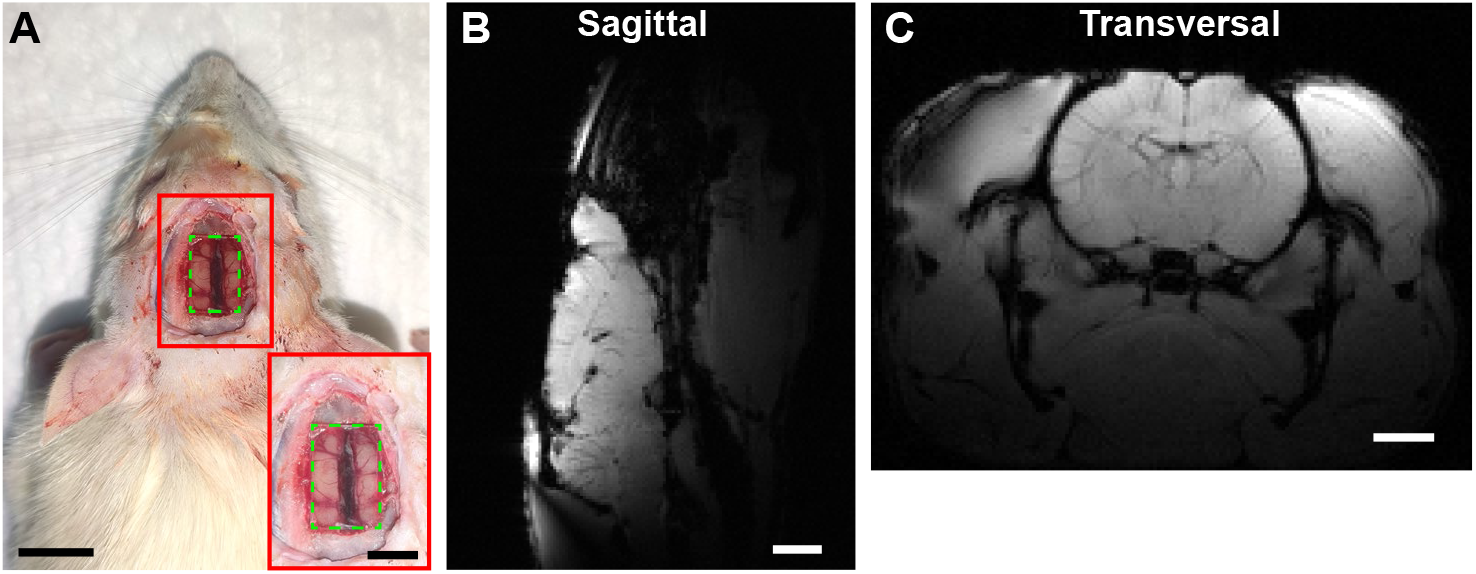
7T MRI study of ZnO-TFTs array on a rat cadaveric model. (A) A photograph of the placement of the ZnO-TFTs array on the surface of rat cadaver brain. Scale bar: 5 mm. Inset: a closer view of the implanted ZnO-TFTs array. Scale bar: 5 mm. The green frames label the location of the electrode array. (B) Sagittal T1-weighted MRI image of a rat brain. Scale bar: 2 mm. (C) Transversal T1-weighted MRI image of rat brain. Scale bar: 2 mm.

## Conclusion

An active transparent electrode array for electrocorticogram electrical recording was proposed, which used ZnO-TFTs as local neural signal amplifier. The employment of ITO and ZnO achieved the most transparent active ECoG array to date (up to 85%), and enabled in-situ and synchronous neural recording and light stimulation for optogenetics on mice brain.

A ZnO-TFTs fabrication technology compatible with that for the electrode array was developed. To enhance the amplification capability of the ZnO-TFTs, the parameters of ZnO-TFTs were optimized by both SPICE model simulation and fabrication. The electrical, optical and mechanical properties of the ZnO-TFTs electrode array were characterized. To further verify the adaptability of our electrode array in recording electrocorticography signal, a series of characterizations were taken to prove its constant level of gain in varied frequency, extremely low latency time, and consistence in electrical properties. The ability for longterm storage was up to 6 months, and waterproof performance of a 20-nm-thick Al2O3 encapsulation was also verified.

The ZnO-TFTs electrode array was applied on a rat brain for sleep waveform and on a mice brain for optogenetics. The former verified its feasibility of electrocorticography signal recording, and the latter proved its high transparency allowing for light stimulation in optogenetics. The SNR was enhanced by the adoption of pre-amplifier, reaching up to 19.9 dB, which was superior to the passive electrodes (Au). Moreover, the array did not blur MRI imaging at 7T ultra-high magnetic field, demonstrated its great potential in MRI-ECoG synchronous recording. In addition, its high transparency can be compatible with techniques of optical brain imaging, which can be applied to more studies combining optics and electrophysiology in the future. As a multimodal active ECoG array, it provides a promising tool for monitoring electrophysiological signals with synchronized optical modulation, 7T MRI and opical imaging.

## Acknowledgement

This work was sponsored by the China Brain Project (2021ZD0200401 and 2022ZD0208600). This project was also supported by Zhejiang Province Key Research & Development programs under the grant of 2021C03003, 2021C03062 and 2021C03108. We also acknowledge the Micro/Nano Fabrication Center, International Campus, Zhejiang University. We would like to thank Prof. Xiaotong Zhang (College of Electrical Engineering, and the Interdisciplinary Institute of Neuroscience and Technology at the School of Medicine, Zhejiang University) for technical supports of 7T MRI.

## Supporting Information

### Multi-layer process

A 4-inch highly-transparent Corning Eagle X Glass (500 µm of thickness) was employed as the substrate. The first ITO layer (In_2_O_3_ : SnO_2_ = 90wt%: 10wt%) of 100 nm thickness was sputtered and patterned by photolithograph as source and drain of the transistors. ITO was then treated with rapid thermal annealing (RTA) at 400 °C for 5 min in nitrogen environment to gain high transparency and electrical conductivity. A ZnO active layer of 20 nm thickness was deposited by thermal atomic layer deposition (ALD, Lesker 150LX) at 200 °C and then annealed at 400 °C for 5 min in oxygen ambient. An Al_2_O_3_ layer of 10 nm thickness was deposited by ALD to protect the ZnO layer and then annealed in oxygen at 200 °C for 30 s. Then the ZnO layer was patterned by photolithograph and wet etching to form the conductive channel of the TFTs, and annealed at 400 °C for 5 min in oxygen ambient. The second Al_2_O_3_ layer, as the gate dielectric layer, was deposited by ALD and annealed at 200 °C for 30 s in oxygen environment. Through-holes were patterned and opened by wet etching on the second Al_2_O_3_ layer, and then the sample was annealed at 400 °C for 5 min in nitrogen environment. Then, the gate electrodes and bonding pads of a 100-nm ITO layer were formed by photolithography and lift-off process via sputtering, and annealed at 400 °C for 5 min in oxygen ambient. Deuterium ions was injected into the transistors by plasma enhanced chemical vapor deposition (70 W, 15 min) to adjust threshold voltage of the ZnO-TFTs. Finally, the third Al_2_O_3_ layer of 20 nm thickness was deposited on the whole device except the gate electrodes and bonding pad region, which was used as the protective layer of the whole device to prevent the entry of moisture and oxygen in air to penetrate into the device that deteriorate the device properties.

### Connection

The wafer was cut into rectangular pieces for each ZnO-TFTs array by a Computer Numerical Control glass cutting machine. The ZnO-TFTs array was bonded with flexible printed circuit (FPC) by anisotropic conductive film (ACF, AC-2056R-35, Hitachi) using a hot press machine (170°C, 0.25 MPa lasting 10 s). Then, the epoxy resin AB glue was applied on the attachment area to avoid breakage of connection as well as the potential risk of electrical leakage caused by the exposed contact.

**Fig. S1.**
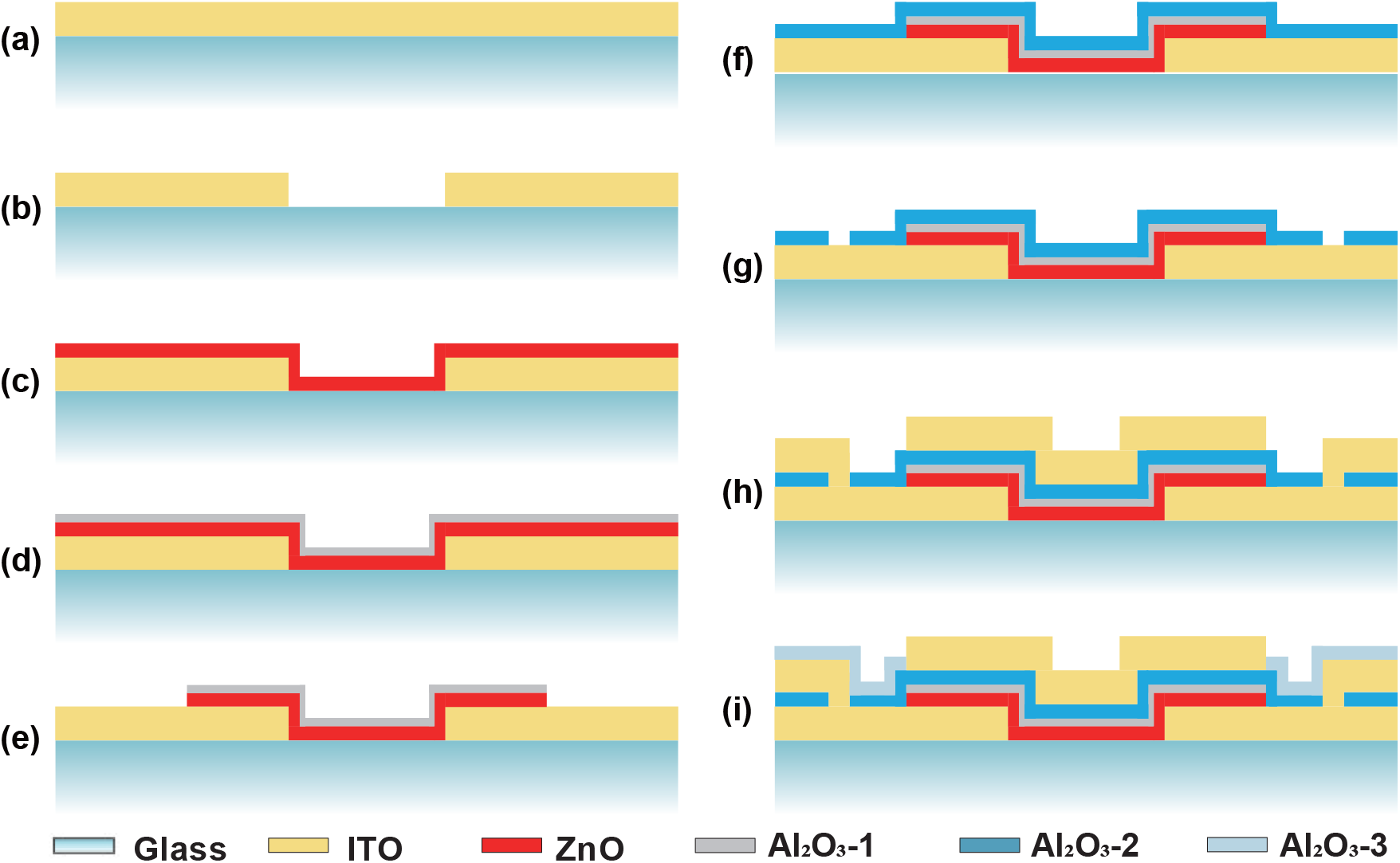
Fabrication process of ZnO-TFTs electrodes array.

**Figure S2.**
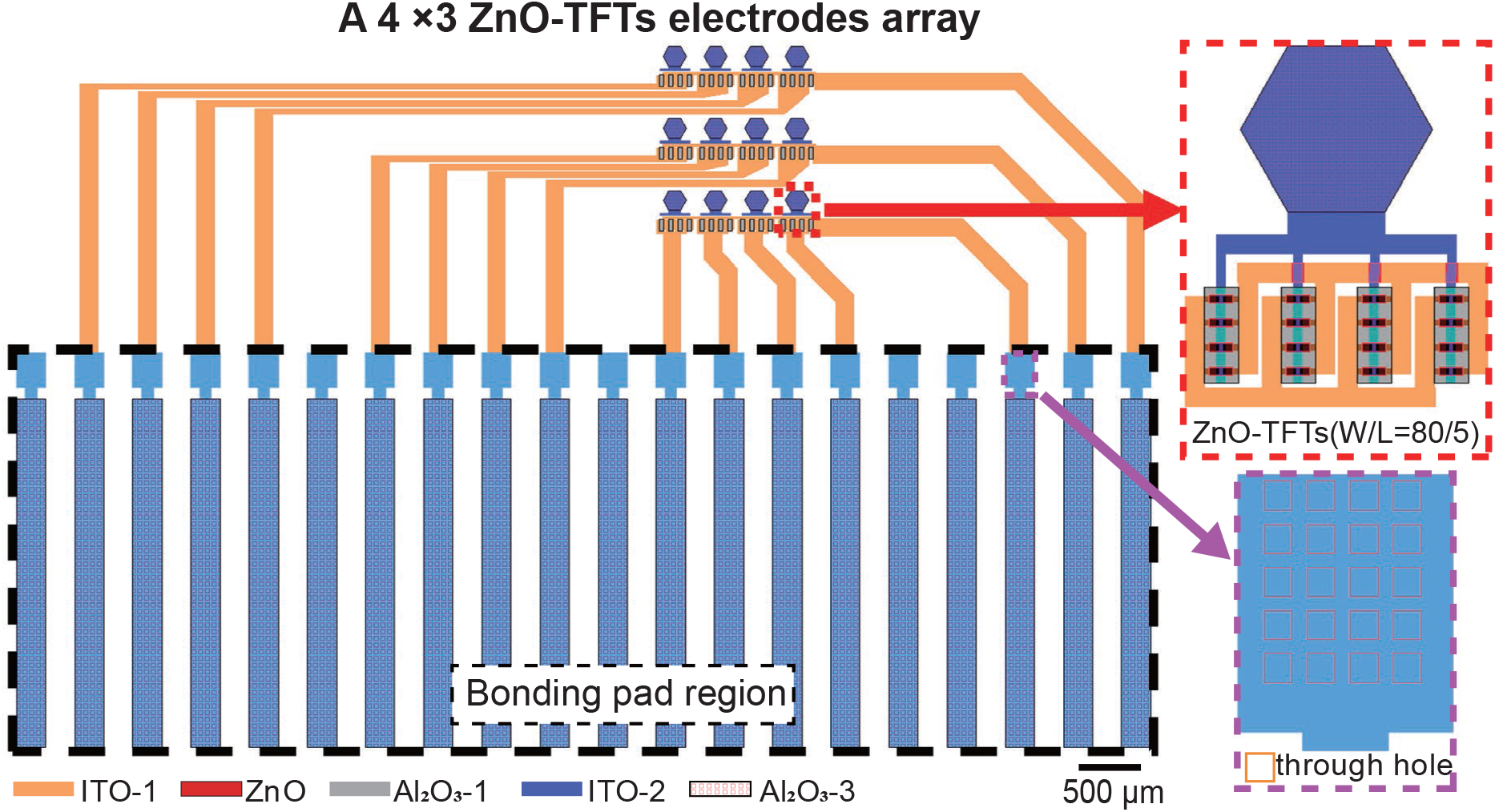
(A) The photograph of ZnO-TFTs measurement on the probe station system. (B) Microscopic image of ZnO-TFTs (W/L=80/5) measurement on the probe station system. (C) Microscopic image of 3 *×* 4 ZnO-TFTs array measurement on the probe station system.

**Figure S3.**
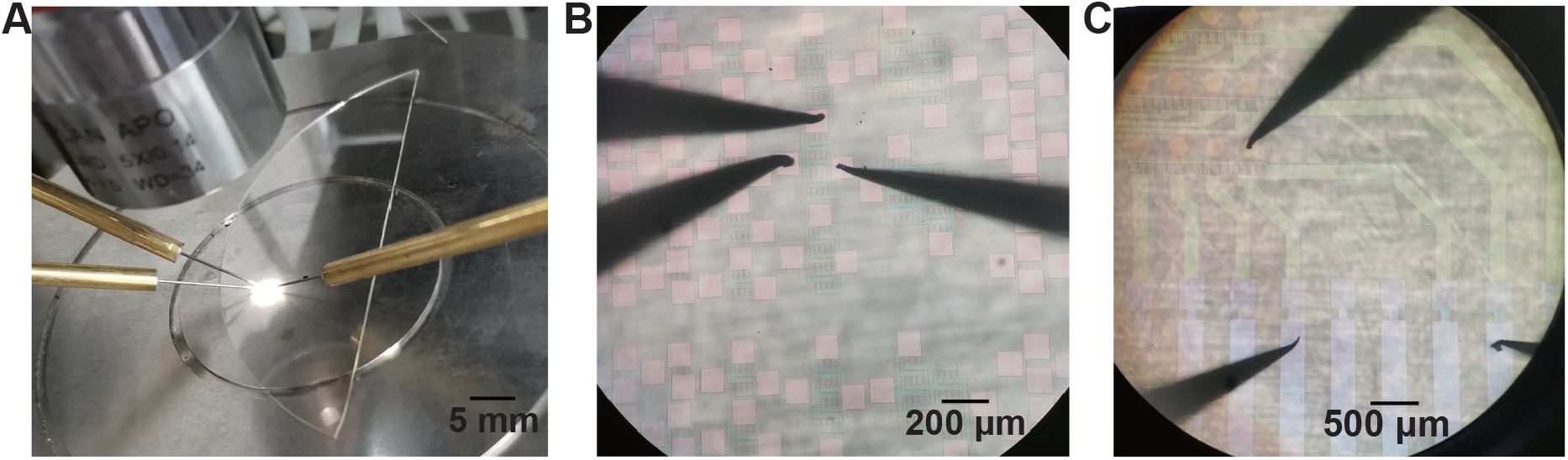
Layout of a 3 × 4 ZnO-TFTs electrodes array. The second Al_2_O_3_ layer covered the whole device except of the through holes.

**Figure S4.**
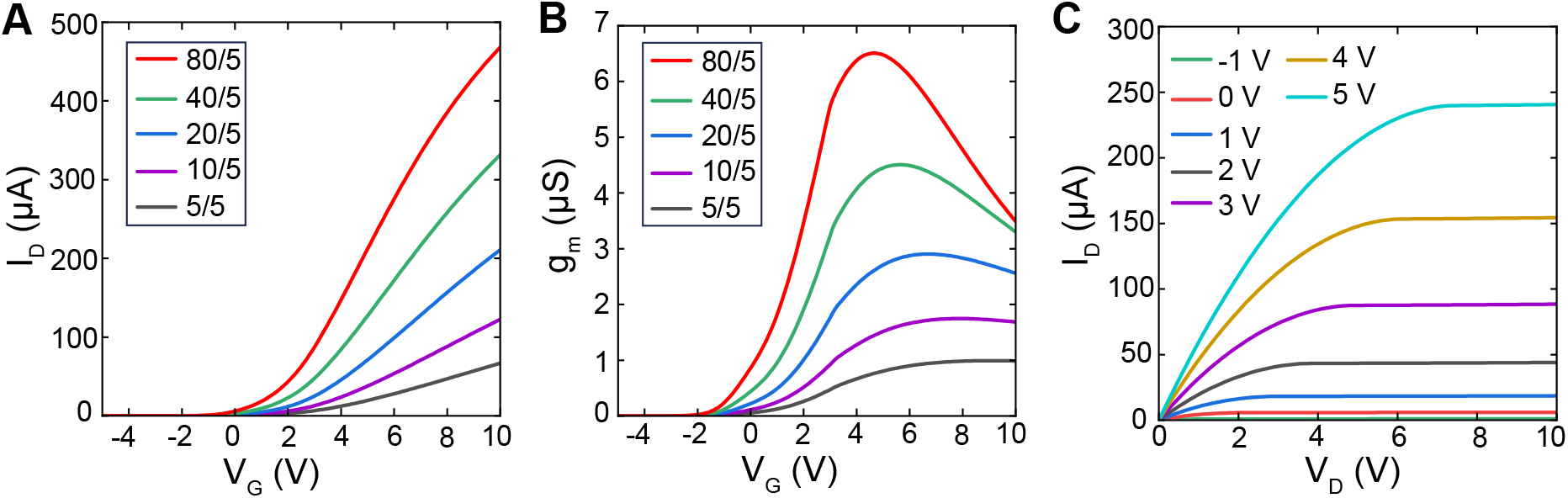
Electrical performance simulation for ZnO-TFTs through n-type LTPS-TFT SPICE model (level 62). (A) The transfer characteristic curves simulated by ZnO-TFTs with various width to length ratio (W/L). (B) The transconductance curves calculated from the transfer characteristics of panel A. (C) The output characteristic curves simulated by ZnO-TFTs with W/L=80/5, with drain voltage, (*V*_*D*_) with gate voltage (*V*_*G*_) varying from -1 V to 5 V (step = 1 V).

**Figure S5.**
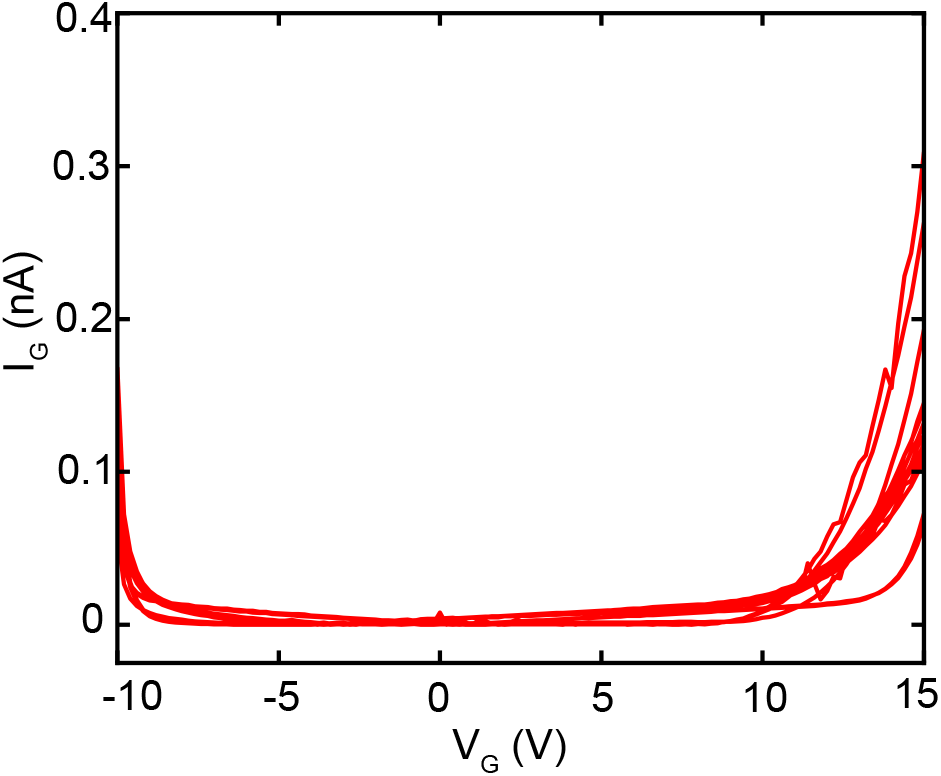
The steady state gate current measured with *V*_*G*_ between -10 V and 15 V, and *V*_*D*_ of 5 V.

**Figure S6.**
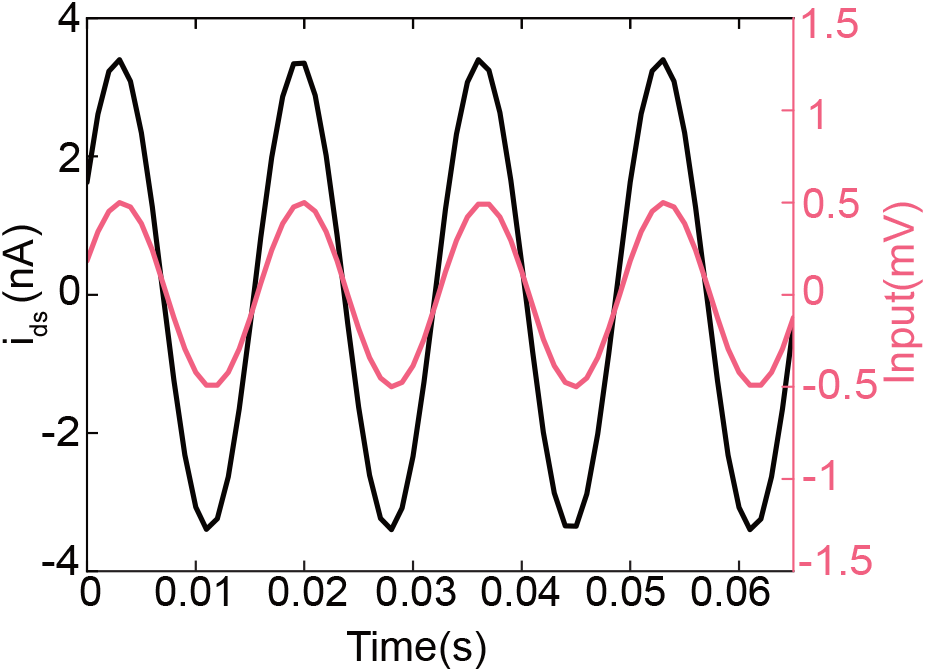
The sine wave with frequency of 60 Hz acquired by ZnO-TFTs array. Input: 60 Hz, peak to peak voltage (*V*_*p−p*_) of 1 mV, sine wave.

**Figure S7.**
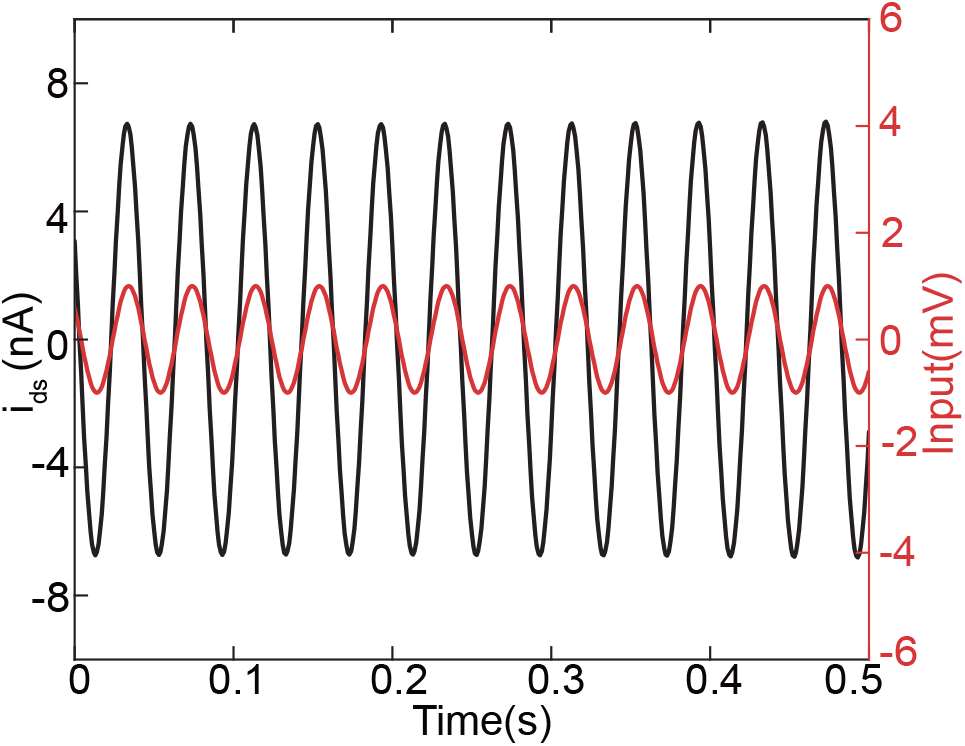
The 25 Hz sine wave acquired by the ZnO-TFTs electrode that has been soaked in saline solution one week ago. Input: 25 Hz, peak to peak voltage (*V*_*p−p*_) of 2 mV, sine wave.

**Figure S8.**
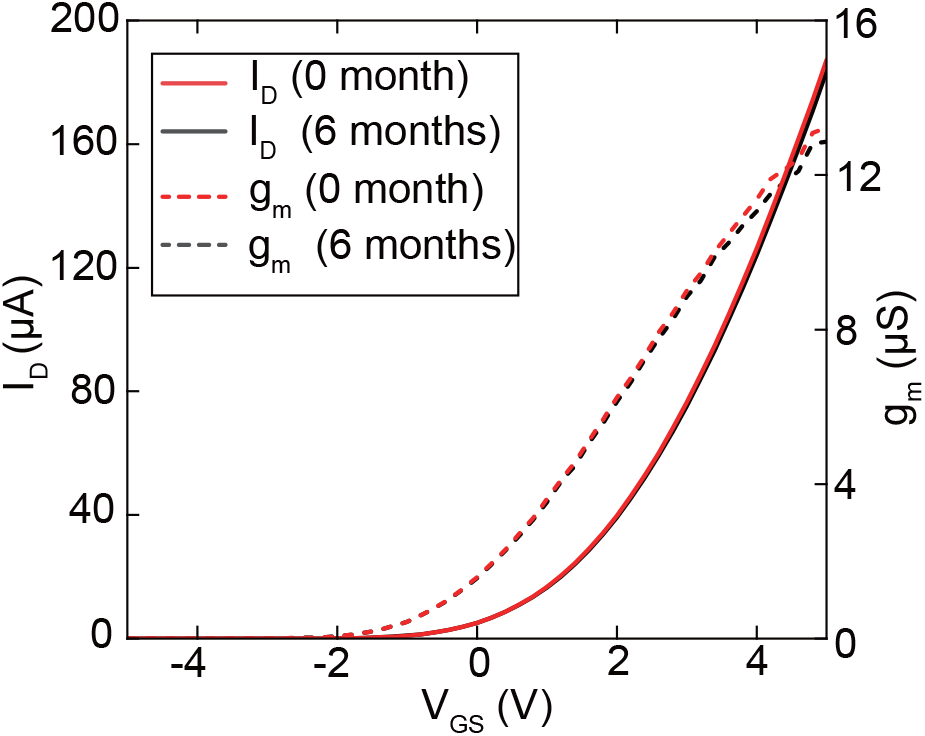
Electrical properties of a ZnO-TFTs electrode stored in dry environment for 6 months, demonstrating the electrical stability of ZnO-TFTs electrodes.

**Figure S9.**
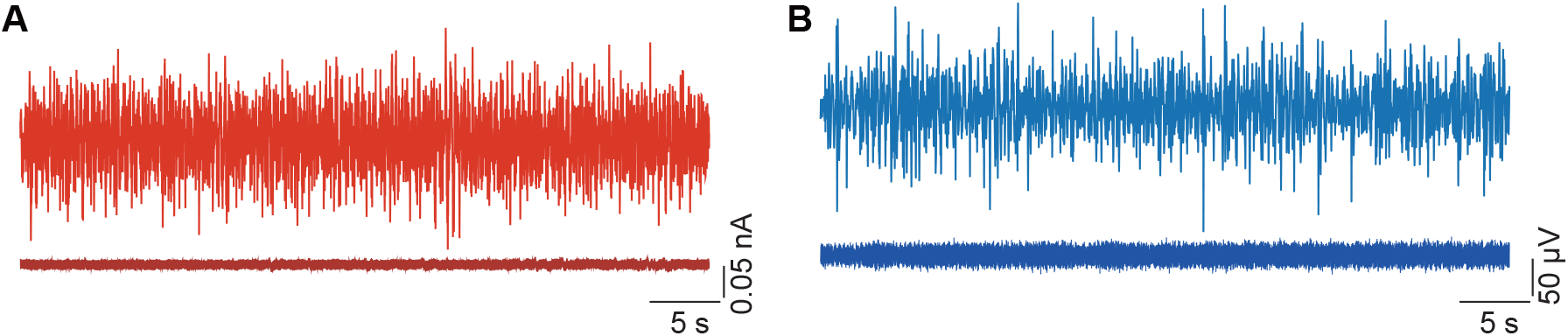
(A) Electrophysiological signal from one ZnO-TFTs electrode (red) and noise recorded on died rat brain (deep red). (B) Electrophysiological recording from one Au electrode (blue) and noise recorded on died rat brain (deep blue).

**Figure S10.**
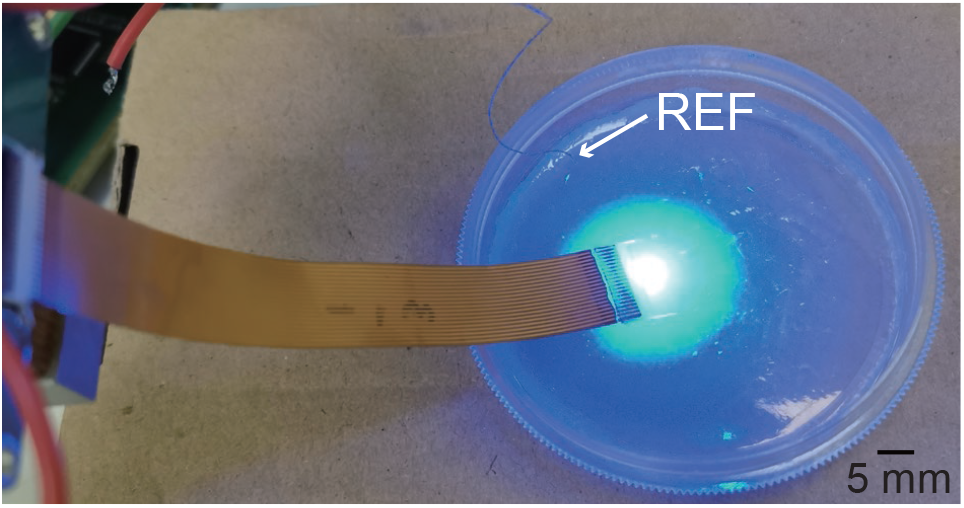
Light-induced artifacts evaluation by positing ZnO-TFTs electrodes array on agar and applying continuous light pulse stimulation.

**Figure S11.**
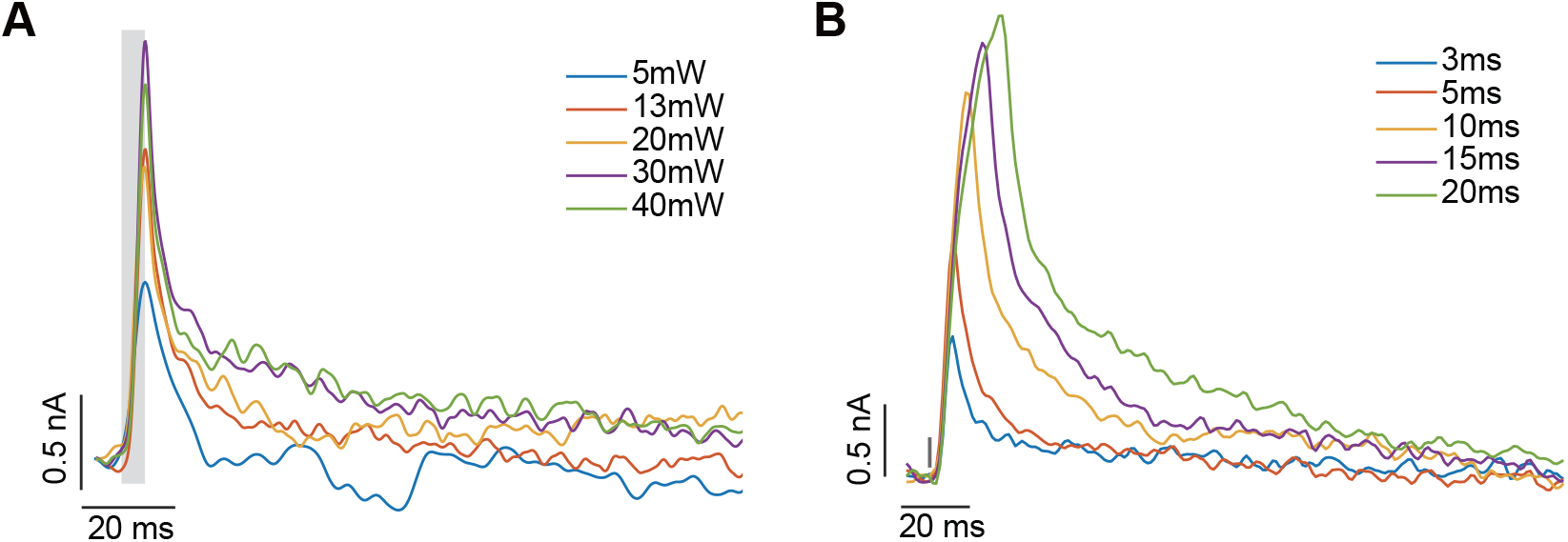
Averaged light-induced artifacts of ZnO-TFTs electrodes array. (A) Averaged light-induced artifacts (n = 30) under light stimulation with various power (4 Hz, 5 ms). The gray rectangles illustrated the start and duration of light stimulation pulses. (B) Averaged light-induced artifacts (n = 30) under light stimulation with different pulse width (4 Hz, 13 mW). The gray line labled the start of light stimulation pulse.

**Figure S12.**
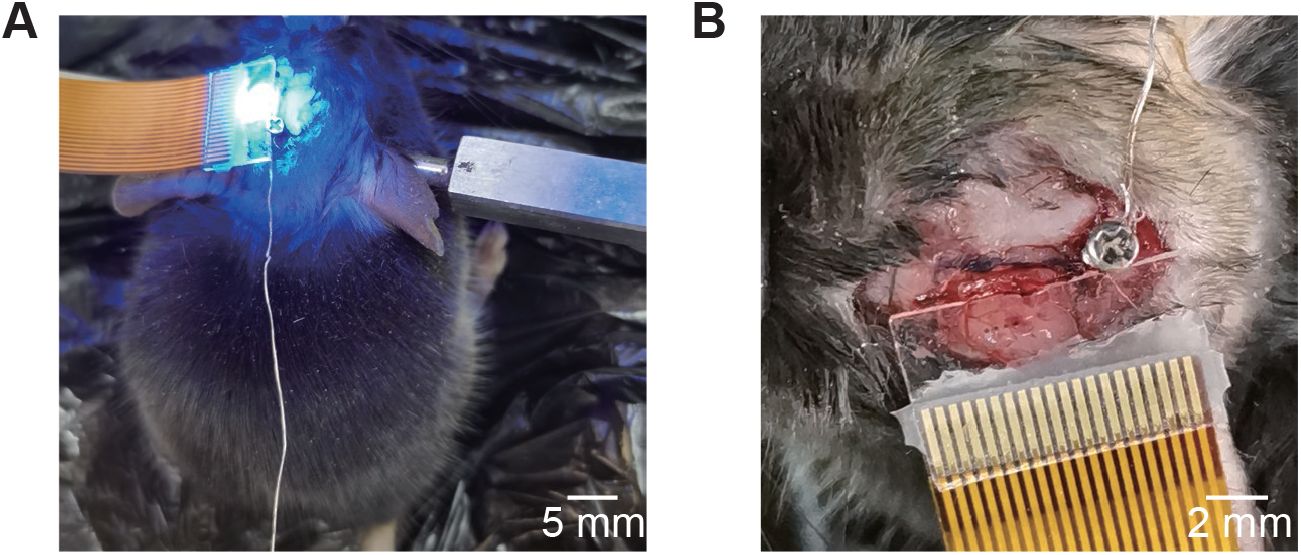
(A) The image of optogenetic experiment using ZnO-TFTs electrodes array. (B) The ZnO-TFTs electrodes array posited on the brain surface of a mice.

**Figure S13.**
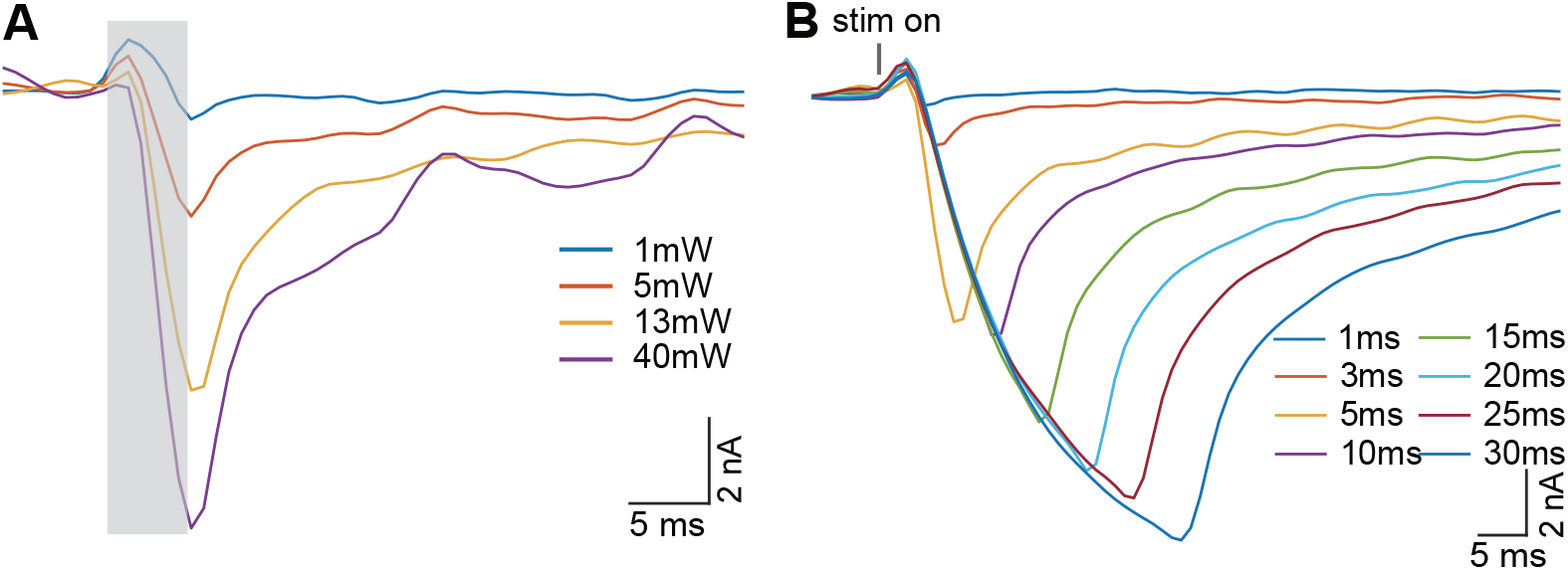
Averaged light-evoked potential (n = 30) recorded by ZnO-TFTs electrode under photostimulus with different light power (A) and duration (B). In (A), 5 ms, 4 Hz light pulse trains were applied, and the gray rectangles illustrated the start and duration of light stimulation pulses. In (B), 13 mW, 4 Hz light pulse trains were applied. The gray line labled the start of light stimulation pulse.

